# Air pollution mediates diabetes by disrupting gut innate mucosal immunity

**DOI:** 10.1101/827014

**Authors:** Angela J. T. Bosch, Theresa V. Rohm, Shefaa AlAsfoor, Zora Baumann, Marc Stawiski, Thomas Dervos, Sandra Mitrovic, Julien Roux, Daniel T. Meier, Claudia Cavelti-Weder

## Abstract

Air pollution has emerged as an unexpected risk factor for diabetes^1–6^. The mechanisms linking air pollution and diabetes remain, however, unknown. It has been postulated that lung exposure mediates diabetes via systemic inflammation and insulin resistance^7–9^. By contrast, gut exposure to pollutants has received little attention, even though a large proportion of air pollution particles are swallowed after mucociliary clearance from the upper airways^10^. Here, we identified intestinal macrophages as key mediators linking air pollutants with impaired beta-cell function. Upon oral exposure to air pollution, intestinal macrophages up-regulated inflammatory- and interferon-response pathways, which disrupted their normal differentiation towards an anti-inflammatory/resident phenotype. The resulting pro-inflammatory milieu in the gut impaired beta-cell identity and function via local cytokine secretion from macrophages as genetic and pharmacological macrophage or IL-1β ablation protected mice from developing air pollution-induced diabetes. These data establish intestinal macrophage-derived cytokines as key mediators of air pollution associated beta-cell dysfunction, thus pointing towards novel therapeutic strategies.

---

Air pollution has become one of the most urgent health issues worldwide, causing 6.5 million premature deaths in 2015^11^. Additionally, air pollution has emerged as an unexpected risk factor for diabetes in many epidemiological^1–6^ and rodent studies^12,13^. This association even occurs at air pollution levels well below those labelled as safe by the World Health Organization^14^. However, underlying mechanisms remain poorly understood, especially as air pollution particles do not seem to reach the systemic circulation^15^. To date, the lung has been considered as the main target organ of air pollution. By contrast, the gut has not been considered, despite the fact that the majority of pollution particles eventually reach the gut by mucociliary clearance from the airways^10^. Pollutants contaminate food and water, accounting for additional sources of oral exposure^16^. Underlining the clinical relevance of gut exposure, air pollution has been linked to increased incidence of many gastrointestinal tract diseases, such as inflammatory bowel disease, appendicitis, and irritable bowel syndrome^17^. Moreover, air pollutants are known to alter gut microbiota^18,19^, increase gut leakiness^20^, and induce gut inflammation^21^. Thus, the effects of air pollution on glucose metabolism could be caused by gut exposure.

First, we evaluated the effect of exclusive lung exposure on glucose metabolism by exposing mice on standard diet or on combined high-fat diet/streptozotocin (HFD/STZ; type 2 diabetes model) to diesel exhaust particles (DEP), particulate matter (PM), or PBS (phosphate-buffered saline; control) by intratracheal instillation (30 μg twice weekly). Glucose tolerance of HFD/STZ mice intratracheally exposed to DEP or PM for 3 months did not differ from the controls exposed to PBS. There were no changes in body weight, insulin, and fasting glucose (Fig. 1a). Accordingly, mice on standard diet who were intratracheally exposed to DEP or PM for 6 months did not develop impaired glucose tolerance and had no changes in insulin, body weight and fasting glucose (Fig. 1b). Macroscopically, lungs from intratracheally exposed mice appeared black, confirming that pollution particles reached the lungs (Fig. 1c). These deposits caused lung inflammation, as shown by increased frequencies of monocyte-derived CD11b^+^ macrophages and eosinophils following intratracheal exposure, while frequencies of tissue-resident alveolar macrophages and conventional dendritic cells (DCs) were unchanged (Fig. 1d, Extended Data Fig. 1). Lung exposure did not elicit systemic inflammation (TNF-α, IL-6), but blood cholesterol was elevated (Fig. 1e,f). Hypercholesterinemia was mirrored by increased liver lipids, while liver enzymes and inflammatory gene expression were mostly unchanged (Fig. 1g, Extended Data Fig. 1). Adipose tissue inflammation, the hallmark of tissue inflammation in metabolic disease, did not develop (Fig. 1h). Thus, although exclusive lung exposure to pollutants causes lung inflammation in mice, it does not result in systemic inflammation and diabetes.

**Fig. 1.**
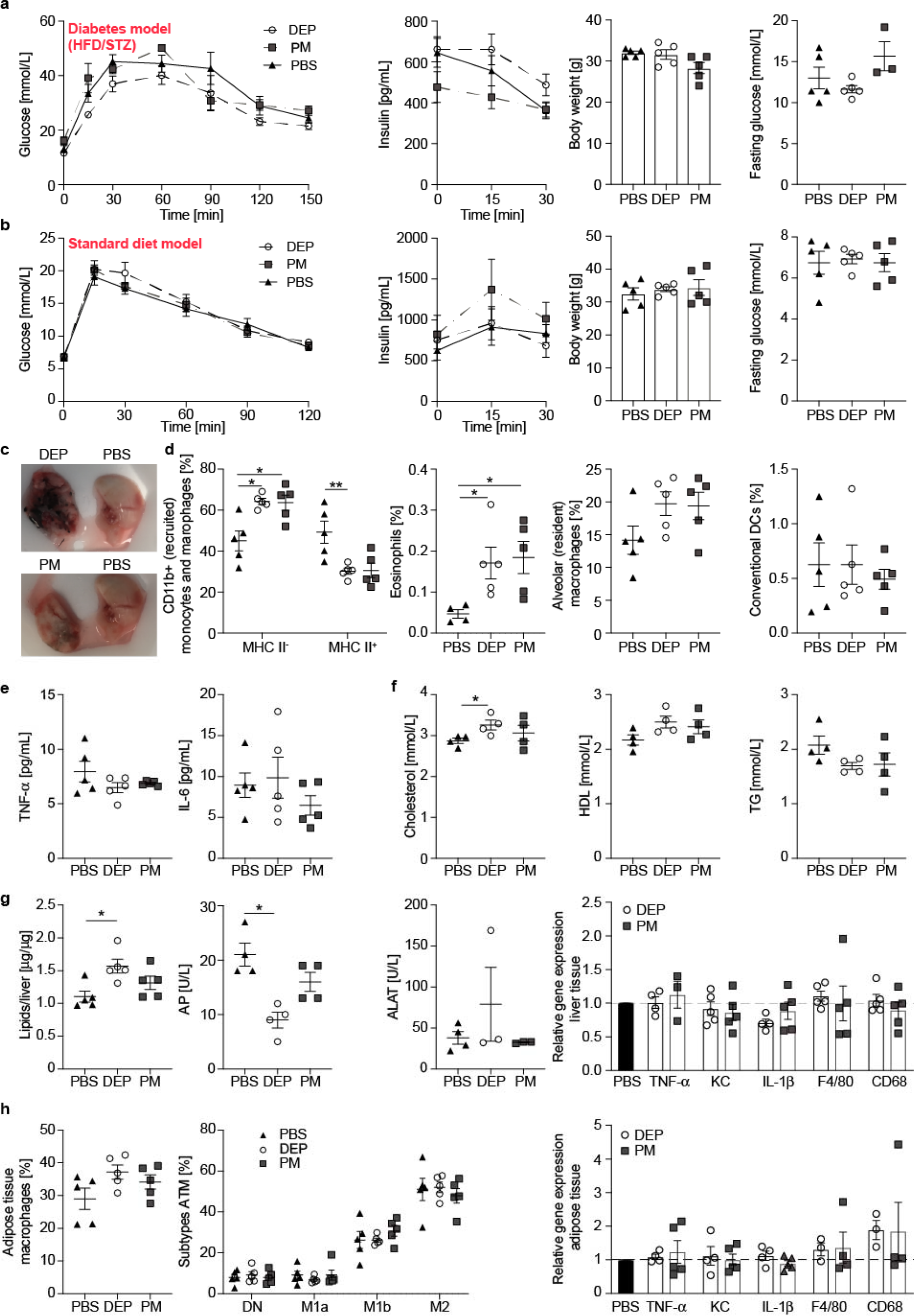
Exclusive lung exposure to air pollutants induces lung inflammation, but not diabetes. Glucose tolerance test (GTT), insulin, body weight and fasting glucose in HFD/STZ wild-type mice after 3 months (**a**) and mice on standard diet after 6 months of intratracheal exposure to DEP, PM, or PBS (**b**). (**c**) Representative pictures of lungs intratracheally exposed to DEP, PM, or PBS. (**d**) Frequencies of lung monocytes and macrophages among CD11b^+^ cells (MHC II^−^ cells correspond to monocytes and macrophages; MHC II^+^ cells to CD11b^+^ DCs and interstitial macrophages), eosinophils (SiglecF^+^CD11c^+^), alveolar macrophages (SiglecF^+^CD11b^−^) and conventional DCs (CD103^+^CD11b^+^) gated on CD45^+^lineage^−^ cells. (**e**) Plasma TNF-α) and IL-6. (**f**) Cholesterol, high-density lipoproteins (HDL), and triglycerides (TG). (**g**) Liver lipids, enzymes (alkaline phosphatase (AP), alanine transaminase (ALAT)), and inflammatory gene expression. (**h**) Adipose tissue macrophage (ATM) frequencies (ATM; F4/80^+^ among CD45^+^), subtype distribution according to CD11c and CD206 and gene expression. Data are presented as meanmSEM of 5 mice per group from one representative experiment compared by a two-tailed, unpaired Mann-Whitney U test (*p<0.05, **p<0.01).

Next, we assessed the role of exclusive gut exposure in mediating air pollution-induced diabetes. Mice on a standard diet or HFD/STZ (type 2 diabetes model) were exposed to DEP, PM, or PBS by oral gavage (12 μg/day on 5 days/week). Importantly, oral and intratracheal exposures were carried out in parallel and contained the same weekly dose of 60 μg, which is equivalent to an inhalational exposure of approximately 160 μg/m^3^. HFD/STZ mice that were orally exposed to air pollutants developed worsened glucose tolerance and decreased insulin secretion from 5 weeks of exposure onwards compared to controls. The body weights of mice exposed to DEP were reduced, reflecting the loss of insulin’s anabolic function. Fasting glucose and insulin sensitivity were unchanged (Fig. 2a). The mice on standard diet exposed to oral air pollution also exhibited impaired glucose tolerance and reduced insulin from 2-4 months onwards, while their body weight, fasting glucose, and insulin sensitivity remained unchanged (Fig. 2b). Exposure to DEP induced more pronounced effects on glycemia than PM; however, there was no dose-dependency when comparing 12 and 60 μg DEP daily (Extended Data Fig. 2). Systemic TNF-α, IL-6 and lipids were not altered, but liver lipids increased similar to mice with lung exposure (Fig. 2c-e, Extended Data Fig. 3). Neither adipose tissue, liver, nor lung inflammation occurred with oral exposure (Fig. 2e,f, Extended Data Fig. 3). Thus, exclusive gut exposure to air pollution leads to diabetes in mice due to reduced insulin secretion, but not as a consequence of systemic inflammation and insulin resistance.

**Fig. 2.**
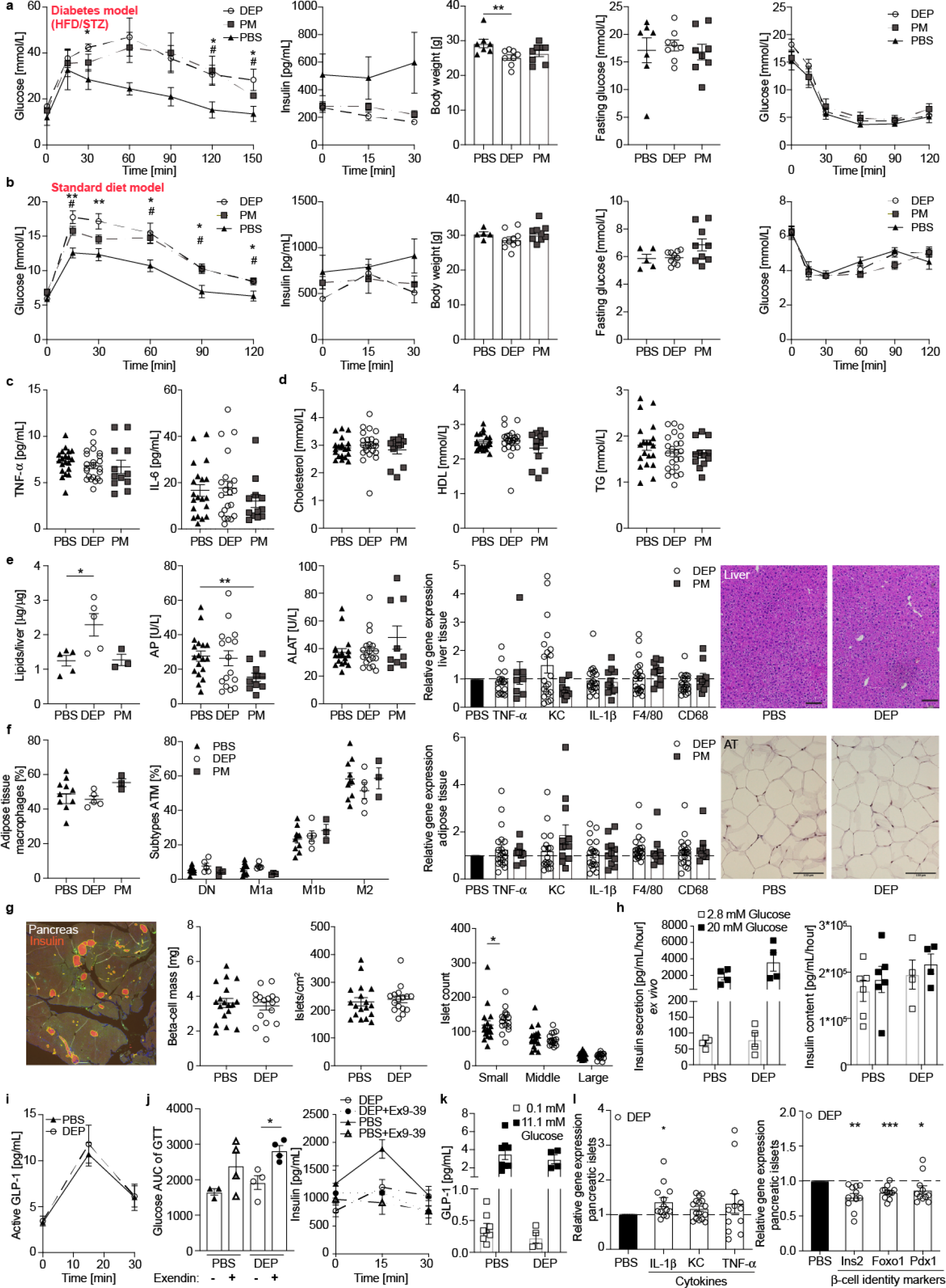
Exclusive gut exposure to air pollutants induces diabetes due to defective insulin secretion. GTT, insulin, body weight, fasting glucose and insulin tolerance test (ITT) in HFD/STZ wild-type mice (**a**) and mice on standard diet after 3 months of oral exposure to DEP, PM, or PBS (**b**). (**c**) Plasma TNF-α) and IL-6. (**d**) Cholesterol, HDL, and TG. (**e**) Liver lipids, enzymes, inflammatory gene expression and H&E staining. (**f**) ATM frequencies (F4/80^+^ among CD45^+^), distribution of subtypes according to CD11c and CD206, gene expression and H&E stainings. (**g**) Insulin staining in pancreatic tissue for analysis of beta-cell mass, number of islets, and size distribution (small 10-1,000μm^2^, middle 1,000-10,000μm^2^, large 10,000-100,000μm^2^). (**h**) *Ex vivo* glucose-stimulated insulin secretion and insulin content of isolated islets from exposed mice. (**i**) Active GLP-1 in oral GTT. (**j**) Glucose AUC of GTT and insulin after GLP-1 antagonist exendin(9-39) injection. (**k**) GLP-1 secretion of primary colon cultures treated *ex vivo* with DEP or PBS. (**l**) Gene expression in islets from exposed mice relative to controls. a/b/f/h depict a representative from 2-5 independent experiments of 5-10 mice. c/d/e/g/l are meannSEM pooled from 3 independent experiments and i/j from one experiment. *p<0.05, **p<0.01, ***p<0.001, unpaired Mann-Whitney U test with two tailed distribution.

We then investigated whether reduced insulin secretion is due to decreased numbers of beta-cells or a functional beta-cell defect. Beta-cell mass was not reduced and the number of small islets even slightly increased in mice exposed to DEP (Fig. 2g). While insulin secretion was impaired *in vivo* (Fig. 2a,b), glucose-stimulated insulin secretion from islets isolated from mice exposed to DEP or PBS was comparable *ex vivo* (Fig. 2h). Reduced insulin secretion was not due to impaired glucagon-like peptide-1 (GLP-1) secretion by enteroendocrine cells as we found similar amounts of active GLP-1 upon oral glucose stimulation and comparable glucose tolerance upon blocking GLP-1 by its receptor antagonist exendin(9-39) in mice exposed to DEP and controls (Fig. 2i,j). Further, primary colon cultures treated with or without DEP *ex vivo* showed similar GLP-1 secretion (Fig. 2k). When directly quantifying gene expression of islets from mice exposed to DEP or PBS, we found up-regulation of the pro-inflammatory cytokine *IL-1β*, while beta-cell identity markers *Ins2*, *Foxo1*, and *Pdx1* were reduced (Fig. 2l). Thus, the insulin secretion defect upon DEP *in vivo* is not due to reduced beta-cell mass, but rather due to altered beta-cell identity and function, potentially through enhanced IL-1β secretion.

To examine the role of gut inflammation in inducing this altered islet cell phenotype, we first characterized adaptive immune cells of the gut. In wild-type mice exposed to DEP, intraepithelial lymphocytes and their subsets were unaffected (Fig. 3a). Also, lamina propria T-cells, dendritic cells, and innate lymphoid cells were unchanged (Fig. 3b-d). To conclusively assess whether the adaptive immune system is required to mediate air pollution-induced diabetes, we exposed Rag2−/− mice who lack adaptive immunity to DEP or PBS. The Rag2−/− mice exposed to DEP developed a drastic impairment of glucose tolerance already at one month of exposure (Fig. 3e). While oral DEP led to a loss of CCR2^−^ (“M2”-like) anti-inflammatory/resident intestinal macrophages indicating a change in innate immunity in the gut, there were no signs of systemic or adipose tissue inflammation (Fig. 3e-i). In sum, adaptive immunity and innate lymphoid cells are not involved in air pollution-induced diabetes.

**Fig. 3.**
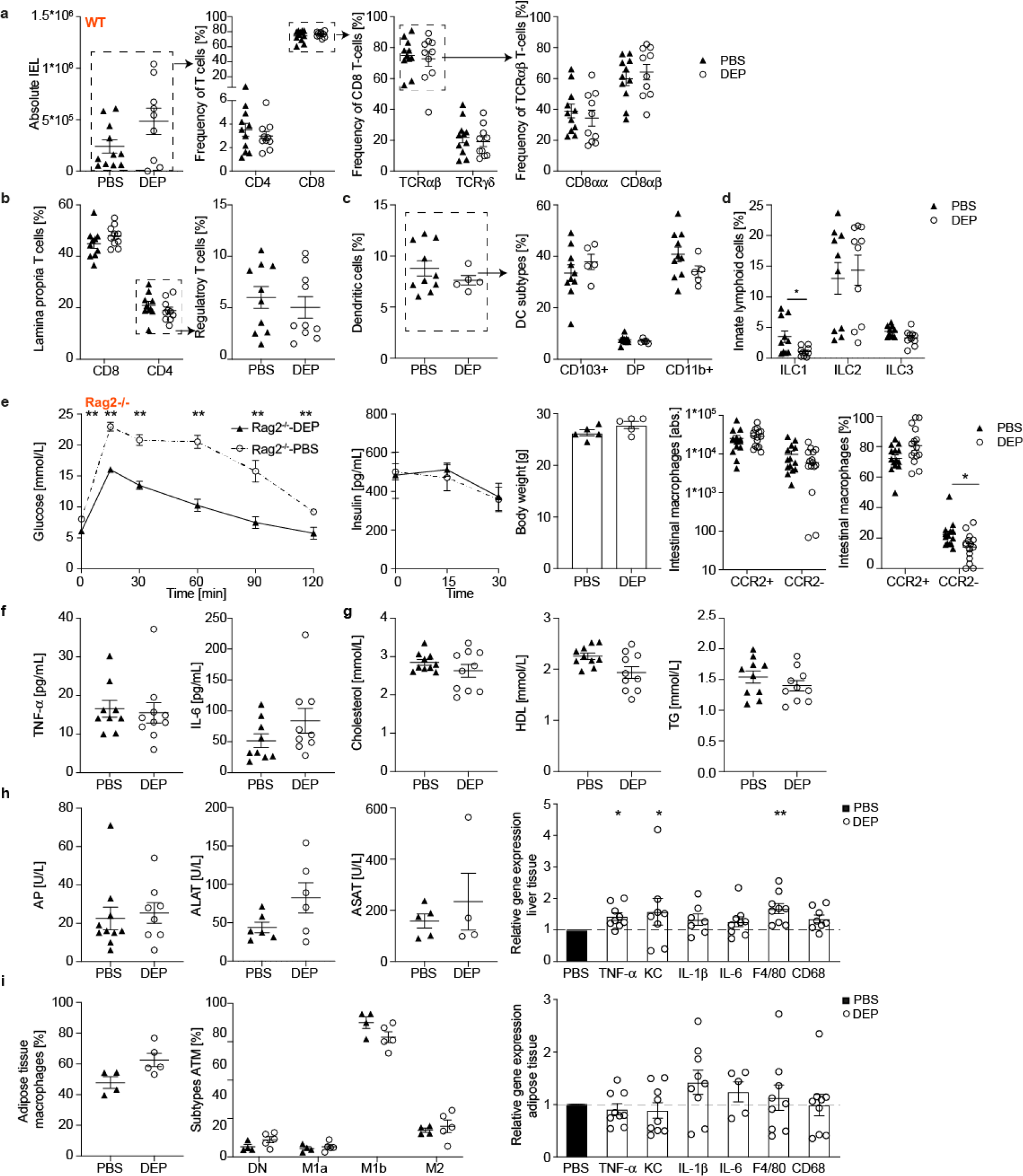
Gut adaptive immunity and innate lymphoid cells are not involved in air pollution-induced diabetes. (**a**-**d**) Colonic intraepithelial and lamina propria cells of wild-type mice orally exposed to DEP or PBS for 6 months (gating strategies see Extended Data Fig. 9): (**a**) Frequencies of intraepithelial cells (CD45^+^CD3^+^) and their subtypes according to CD4, CD8, TCRαβ, TCRγδ, CD8αα, and CD8αβ. (**b**) Frequencies of lamina propria CD4 and CD8 T-cells and regulatory T-cells (Foxp3^+^CD25^+^). (**c**) Frequencies of dendritic cells (CD64^−^MHCII^+^CD11c^+^) and their subtypes according to CD103 and CD11b. (**d**) Frequencies of innate lymphoid cells ILC1 (Nkp46^+^), ILC2 (GATA3^+^), and ILC3 (RORγ^+^). (**e-i**) Oral exposure of Rag2−/− mice to DEP or PBS for 4 months: (**e**) GTT, insulin, body weight, absolute numbers and frequencies of CCR2^+^ inflammatory and CCR2^−^ anti-inflammatory/resident intestinal macrophages. (**f**) Plasma TNF-α) and IL-6. (**g**) Cholesterol, HDL, and TG. (**h**) Liver enzymes and inflammatory gene expression. (**i**) ATM frequencies (F4/80^+^ among CD45^+^), distribution of subtypes and inflammatory gene expression. Data are shown as mean±SEM of pooled data from 2 independent experiments. *p<0.05, **p<0.01, unpaired Mann-Whitney U test with two tailed distribution.

Subsequently, we characterized innate immune cells of the gut as potential mediators of the observed islet cell phenotype. In wild-type mice exposed to DEP, a distinct loss of CCR2^−^ (“M2”-like) anti-inflammatory/resident macrophages was observed, resulting in a relative increase in CCR2^+^ (“M1”-like) inflammatory macrophages (Fig. 4a,b). Consistent with an inflammatory milieu in the gut invoked by DEP, gene expression of colon tissue showed up-regulation of pro-inflammatory cytokines (*TNF-α*, *KC*) and macrophage markers (*CD68*, *Ly6C*, Fig. 4c). To better understand the transcriptional response to DEP, we performed single-cell RNA-sequencing of intestinal macrophages. By using unbiased hierarchical clustering of cells, *Ccr2*^+^ inflammatory and *Ccr2*^−^ anti-inflammatory/resident macrophages were identified as the two main populations (the latter comprising two related clusters, Fig. 4d, Extended Data Fig. 4). First, we corroborated the increase in abundance of *Ccr2*^+^ (“M1”-like) relative to *Ccr2*^−^ (“M2”-like) macrophages upon exposure to DEP (Fig. 4e). Subsequently, we compared the transcriptomes of cells from mice exposed to DEP or PBS, stratifying the analysis by cell clusters to correct for differential abundance. Significantly up-regulated genes upon exposure to DEP were related to macrophage activation and interferon regulation (Fig. 4f, Extended Data Fig. 5). Gene-set enrichment analysis confirmed that in mice exposed to DEP, significantly up-regulated MSigDB Hallmark pathways involved inflammatory, interferon α and α responses, as well as allograft rejection (Fig. 4g, Extended Data Fig. 6). Thus, the normal differentiation from CCR2^+^ (“M1”-like) to CCR2^−^ (“M2”-like) intestinal macrophages is disrupted upon exposure to DEP, leading to a pro-inflammatory milieu in the gut wall associated with the development of diabetes.

**Fig. 4.**
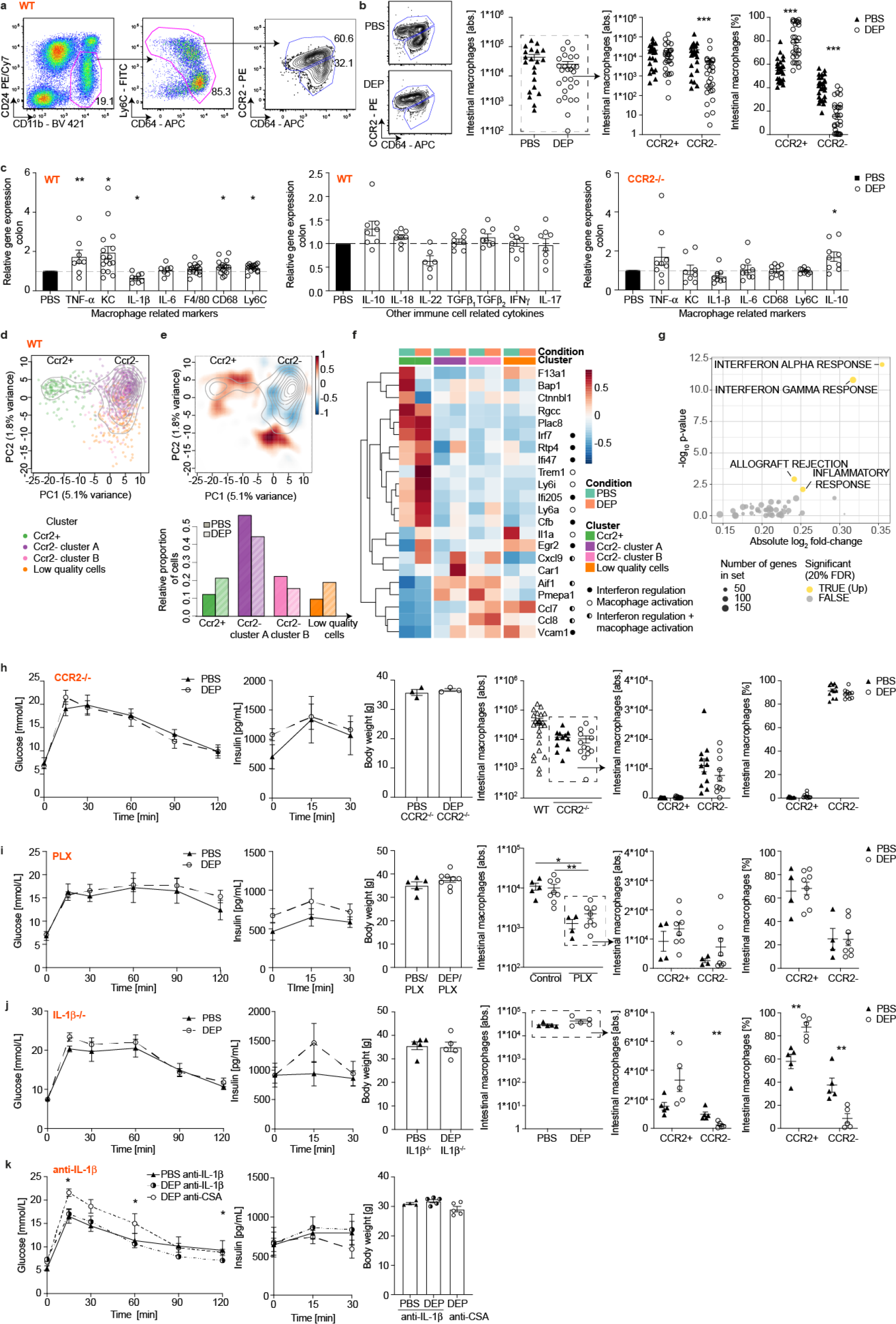
Oral air pollution exposure disrupts intestinal macrophage differentiation, resulting in a predominance of inflammatory macrophages. (**a**) Gating strategy for intestinal macrophages. (**b**) Representative FACS plots, absolute numbers of total, CCR2^+^ inflammatory and CCR2^−^ anti-inflammatory/resident intestinal macrophages and their frequencies in exposed wild-type mice. (**c**) Colon gene expression of immune cell markers in wild-type and CCR2−/− mice exposed to DEP relative to PBS after 5-6 months of exposure. (**d**) Principal component analysis of intestinal macrophages showing *Ccr2*^+^ and *Ccr2*^−^ subpopulations (colors represent different clusters; contour lines indicate cell density). (**e**) Top: Differential 2D density plot (red indicates excess of DEP-, blue excess of PBS-exposed cells); Bottom: relative proportion of cells from mice exposed to DEP or PBS across clusters. (**f**) Cluster-specific differential gene expression of intestinal macrophages in mice exposed to DEP or PBS (closed circles indicate genes related to interferon regulation, open circles genes related to macrophage activation, half-open circles to both). (**g**) MSigDB Hallmark pathways that were significantly enriched in differentially expressed genes upon DEP exposure. (**h-j**) GTT, insulin, body weight, absolute numbers of total, CCR2^+^ inflammatory and CCR2^−^ anti-inflammatory/resident macrophages and their frequencies in exposed CCR2−/− mice (**h**), wild-type mice pharmacologically depleted of macrophages by PLX5622 (**i**), IL-1β−/− mice (**j**), and wild-type mice treated with an anti-IL-1a antibody or a control antibody CSA (anti-cyclosporin A (**k**)). b/c/h/i are mean±SEM pooled from 2-3, and c from 5 independent experiments. d/e/f/g/j/k represent one experiment (for RNA Seq 2 mice per condition). *p<0.05, **p<0.01, ***p<0.001, unpaired Mann-Whitney U test with two tailed distribution.

To establish the causal link between gut innate immunity and air pollution-induced diabetes, we exposed CCR2−/− mice on standard diet to oral air pollution for up to 8 months. CCR2−/− mice lack CCR2-dependent monocyte recruitment and hence CCR2^+^ inflammatory macrophages^22,23^. Accordingly, inflammatory macrophages in the gut of these mice were reduced by 99.5t0.4% and anti-inflammatory macrophages by 57.7a22.4% (Extended Data Fig. 7). In contrast to wild-type mice, there was no shift towards CCR2^+^ (“M1”-like) inflammatory macrophages upon exposure to DEP in CCR2−/− mice, protecting them from a pro-inflammatory milieu in the gut, defective insulin secretion, and diabetes (Fig. 4c,h). As an additional macrophage depletion model, we used a standard diet containing the Colony stimulating factor 1 receptor (CSF1R)-inhibitor PLX5622. CSF1R regulates the survival, proliferation, differentiation, and chemotaxis of macrophages^24^. In mice treated with PLX5622, intestinal macrophages were strongly reduced (inflammatory by 86.7m9.9%, anti-inflammatory by 92.893.8%, Extended Data Fig. 8). When orally exposed to DEP for up to 10 months, these mice retained low numbers of intestinal macrophages, which also prevented the shift towards CCR2^+^ (“M1”-like) inflammatory macrophages, the insulin secretion defect, and diabetes (Fig. 4i). In both models, dyslipidemia, systemic, adipose tissue, and liver inflammation did not occur upon exposure to DEP (Extended Data Fig. 7-8). IL-1β−/− mice that were orally exposed to DEP or PBS for up to 11 months showed a loss in CCR2^−^ (“M2”-like) anti-inflammatory/resident macrophages similar to wild-type mice. However, these mice did not show any impairment in glucose tolerance or an insulin secretion defect, potentially through the loss of macrophage-derived IL-1β secretion (Fig. 4j). Accordingly, treatment with a monoclonal antibody against IL-1β reversed glucose intolerance in mice exposed to diesel (Fig. 4k). Taken together, genetic and pharmacological macrophage or IL-1β ablation prevents or reverts air pollution-induced diabetes. This indicates that intestinal macrophage-derived cytokines are the key mediators of air pollution-induced beta-cell dysfunction.

In sum, long-term exposure of mice to air pollution via gavage leads to impaired glucose tolerance with an inability to mount an adequate insulin response, indicative for an insulin secretion defect. In contrast to our findings, previous studies using whole-body inhalation chambers found impaired insulin sensitivity, potentially as lung and gut exposures occur concurrently^25^. Additionally, most studies were performed in mice on HFD, which per se causes systemic inflammation and insulin resistance, confounding the metabolic phenotype^12,13^. Importantly, only minor amounts of inhaled particles have been detected in the blood circulation^15^, rendering systemic distribution of particles unlikely. We identified intestinal macrophages as the key mediators between air pollutants in the gut lumen and impaired beta-cell function. Indeed, intestinal macrophages can reach into the gut lumen to sample luminal contents^26^. In normal health, intestinal macrophages originate from monocytes and first adopt a CCR2^+^ (“M1”-like) inflammatory phenotype in the gut and then gradually lose their inflammatory phenotype to become CCR2^−^ (“M2”-like) anti-inflammatory/resident macrophages^27,28^. Air pollutants up-regulate inflammatory- and interferon-related pathways in intestinal macrophages, which interferes with their normal differentiation to an anti-inflammatory/resident phenotype. The ensuing pro-inflammatory milieu in the gut then impacts on beta-cell identity and function. Genetic or pharmacological macrophage ablation protects mice from developing a pro-inflammatory milieu in the gut, defective insulin secretion and diabetes. Protection of IL-1β−/− mice and reversal from air pollution-induced diabetes by pharmacological IL-1β blocking suggests a cytokine-driven disease mechanism. The link between inflammation and altered beta-cell identity and function is supported by previous studies: Inflammatory cytokines lead to beta-cell dedifferentiation^29^ and a failure to coordinate insulin secretion within islets^30^, potentially as an adaptive mechanism to escape beta-cell death under stress conditions^31^. Further evidence comes from studies that use anti-inflammatory approaches to improve glucose metabolism^32^. The question remains whether intestinal macrophages directly traffic from the gut to pancreatic islets, which would be against the current dogma, or whether inflammation disseminates to the pancreas through other means such as neuronal circuits, blood circulation or lymphatic vessels. In contrast to mice with defective innate immunity, mice lacking adaptive immunity are not protected and even develop diabetes after shorter exposure to DEP compared to wild-type mice. Their increased susceptibility could be due to enhanced innate immunity as previously reported^33^ and is consistent with our finding of increased ratios of CCR2^+^ inflammatory intestinal macrophages and liver inflammation in Rag2−/− mice exposed to DEP. In sum, our study provides novel insights into how air pollution affects glucose metabolism and represents a crucial advance towards finding better disease prevention and treatment strategies, and thus having a healthier society.

## Methods

### Animals

Male C57BL/6N mice were obtained from Charles River Laboratories (Sulzfeld, Germany). CCR2−/−, IL-1β−/− and Rag2−/− mice were bred in our facility. Mice were maintained in specific pathogen-free conditions with free access to water and standard pelleted food. C57B6/N mice were rendered diabetic by high-fat diet (HFD; Research Diets, New Brunswick, NJ, USA) and a single intraperitoneal (i.p.) dose of streptozocin (STZ; 120mg/kg body weight, Sigma) after 4 weeks of HFD. All procedures were approved by the local Animal Care and Use Committee.

### Exposure protocol

Diesel exhaust particles (DEP; NIST 1650b) or particulate matter (PM; NIST 1649b) dissolved in sterile PBS or PBS alone as control were administered either exclusively to the lung by intratracheal instillation or to the gut by oral gavage for up to 10 months. For intratracheal instillation, mice were anesthetized by using 2% isoflurane. A 22-gauge catheter was inserted into the trachea and the inoculum of 60 μl PBS containing either 30 μg DEP or PM, or PBS alone, passively inhaled into the lungs^34^. This delivery method results in comparable levels of pulmonary distribution and injury biomarkers compared to ambient air exposure^35^. For oral gavage, mice received 12 or 60 μg DEP or PM suspended in 200 μl sterile PBS or PBS as control. Oral gavage was performed 5 days a week, whereas intratracheal instillation was conducted twice a week, adding up to a weekly dosage of 60 μg DEP or PM, respectively.

### Glucose and insulin tolerance tests (GTT/ITT)

GTT and ITT were performed at monthly intervals. Glucose (2g/ kg body weight) was administered either by i.p. injection or oral gavage after 6 hours of fasting. Blood samples were obtained from the tail vein and assayed by using a glucometer (Freestyle; Abott), or mixed with EDTA (Sigma) or diprotein A (Bachem) and spun down for later analysis. To stabilize systemic active GLP-1, 25 mg/kg body weight Sitagliptin (Santa Cruz) was i.p. injected 30 minutes prior to oral glucose administration. To block GLP-1 signaling, synthetic exendin(9-39) 25 nM/kg body weight (Bachem) was i.p. injected 1 minute prior to the GTT. For ITT, mice fasted 3 hours and were injected with 1 U/kg body weight insulin (Novo Nordisk) and glucose measured as specified above.

### Isolation and flow cytometric assessment of immune cells

Cells of the colon, lung and adipose tissue were isolated by enzymatic digestion as follows:

#### Colon

The colon was washed in HBBS (Gibco) and cut into small pieces, placed on an orbital shaker at 37°C and washed twice for 10 minutes in HBSS containing 2mM EDTA (Sigma) and rinsed twice with HBSS prior to digestion with 1 mg/mL collagenase VIII (Sigma-Aldrich) and 12.5 μg/ml DNase I (Sigma-Aldrich) for 30 minutes shaking at 37°C, followed by homogenization with gentleMACS Dissociator (Milteny Biotec) program intestine. Colon cells were enriched for leukocytes using a Percoll gradient (GE Healthcare; 40% and 70%). IEL were isolated from colon tissue by two rounds of 4 minutes vortexing with maximal speed and enriched using Percoll. Single cell suspensions were stained for 30 minutes at 4°C, intracellular staining was performed by using Foxp3 staining kit (eBioscience) according to the manufacturer’s protocol.

#### Lung

Lungs were perfused with PBS, then isolated and minced by using gentleMACS program m_lung_01-02, digested 30 minutes at 37°C on an orbital shaker with 0.15 WünschU/mg Liberase (Sigma) and 0.1 mg/mL DNase I, and homogenized by using gentleMACS program m_lung_02_01. Lung cells were enriched for leukocytes by Percoll as described in the section above.

#### Adipose tissue

Epididymal adipose tissue was minced and digested with 1.5mg/mL collagenase IV (Worthington) and 8.25 μg/mL DNase I for 25 minutes on a thermomixer with 400 rpm. Red cells were removed by red cell lysis buffer.

Adipose tissue macrophages were gated as CD45^+^F4/80^+^CD11b^+^ and subdivided into subpopulations double negative (DN), M1a (CD11c^+^CD206^−^), M1b (CD11c^+^CD206^int^), and M2 (CD11c^− to low^CD206^+^). Cell analysis and sorting were performed on a FACS LSRII Fortessa and FACS Aria III, respectively (BD Biosciences). Acquired data were analyzed using FlowJo software (TreeStar Inc. Ashland, OR, USA). From Biolegend, we obtained antibodies against CD11b (M1/70), CD11c (N418), MHCII (M5/114.14.2), Ly6C (HK1.4), CD45 (30-F11), F4/80 (BM8), CD103 (2E7), CD24 (M1/69), CD64 (X54-5/7.1), CD3 (145-2C11 and 17A2), CD19 (6D5), NK1.1 (PL136), Ly6G (1A8), CD206 (C068C2), CD25 (PC61), CD4 (GK1.5), CD8 (53-6.7), CD8b (YTS156.7.7), TCR-b (H57-597), and TCR-g (GL3). mAb for CCR2 (475301) was purchased from R&D. mAb for Foxp3 (FJK-16s), GATA3 (TWAJ), EOMES (Dan11mag) and Foxp3 staining kit were purchased from Invitrogen. mAb for SiglecF (E50-2440), RORg (Q31-378), NKp46 (gE2/NKp46) were obtained from BD.

### Gene expression analysis

RNA was isolated by using the NucleoSpin RNA kit (Macherey Nagel, Du◻ren, Germany) or the RNeasy Plus Universal Mini kit (QIAGEN, Du◻sseldorf, Germany). Reverse transcription was performed with GoScript™ (Promega). GoTaq qPCR Master Mix (Promega, Madison, WI, USA) was used for real-time PCR (ViiA7, Thermo Fisher Scientific). Primer sequences (Microsynth, Balgach, Switzerland) are listed in Supplementary Table 1.

### Protein expression analysis

Plasma insulin, active GLP-1, TNF-α and IL-6 were quantified by electrochemiluminescence (MESO SECTOR S 600) using kits from MesoScale Diagnostics (MSD, Rockville, MD, USA).

### Liver enzymes and lipids

#### Systemic liver enzymes and lipids

Liver enzymes such as AP, ASAT and ALAT and blood lipids such as cholesterol, HDL and triglycerides were measured in mouse plasma on the c502 / c702 modules of the Cobas 8000 series from Roche Diagnostics (Roche Diagnostics, Basel, Switzerland) according to manufacturer’s instructions.

#### Liver lipids

Approximately 30 mg of liver tissue were homogenized PBS. 2:1 Chloroform-MeOH was added and the samples spun for 5 minutes at 4°C, 3400 rpm. The bottom phase was collected, of which an aliquot was assessed. The samples were first dried, cooled down on ice, after adding H2SO4 boiled at 90°C for 10 minutes and cooled down on ice. After incubation with Vanillin-reagent for 40 minutes, the samples were measured at 550nm on an EPOCH ELISA reader from BioTek instruments.

### Beta-cell mass

For heat-induced antigen retrieval, 5 μm thick sections were boiled for 30 minutes at 93°C in 1x epitope retrieval solution (Biosystems). The slides were stained with the primary antibody for insulin (DAKO) overnight at 4°C, washed twice in PBS (5 minutes), stained with a CD45 antibody (BD Pharmingen) for 2 hours at room temperature and washed twice with PBS (5 minutes). The secondary antibodies (Life Technologies) were applied for 2 hours at room temperature, washed twice with PBS, before mounting with a fluorescence mounting media (Dako). A Nikon microscope at 4× magnification acquired pictures.

Images were analyzed using Fiji software. Analysis was performed in a semi-automated way; ilastik software (https://www.ilastik.org/) was trained twice, once to recognize pancreas area excluding background and lymph nodes, and the second time to recognize islets. The masks ilastik generated were used to quantify the areas with Fiji. Beta-cell mass was defined as area of insulin positive cells/area of pancreas*weight of the pancreas. Two sections per animal were quantified. *Antibodies:* guinea pig-anti-insulin (Dako), rat-anti-CD45 (BD Pharmingen), goat-anti-guineag pig Alexa 657 and goat-anti-rat Alexa 555 (Life Technologies).

### Single-cell RNA-sequencing

Lamina propria cells positive for CD11b and positive either for Ly6C, CD64, or both, were sorted from 2 controls and 2 mice exposed to DEP. Cell suspensions were loaded on the wells of a 10× Genomics Chromium Single Cell Controller (one well per mouse replicate). Single-cell capture and cDNA and library preparation were performed witha Single Cell 3’ v2 Reagent Kit (10× Genomics) according to the manufacturer’s instructions. Sequencing was performed on one flow-cell of an Illumina NexSeq 500 machine at the ETH Zurich Genomics Facility in Basel. Data were analyzed by the Bioinformatics Core Facility, Department of Biomedicine, University of Basel. Paired-end reads were obtained and their quality was assessed with the FastQC tool (version 0.11.5). Sequencing files were processed with the Cell Ranger software (version 3.0.2, provided by 10× Genomics at https://support.10xgenomics.com/single-cell-gene-expression/software/downloads/3.0 to perform sample and cell demultiplexing, read alignment to the mouse mm10 genome assembly with STAR, and to generate read count table. Default settings and parameters were used, except for the version of STAR updated to 2.6.1a^36^, and the STAR parameters *outSAMmultNmax* set to 1 and *alignIntronMax* set to 10,000. The reference transcriptome “refdata-cellranger-mm10-3.0.0”, provided by 10× Genomics and based on Ensembl release 93^37^, was used (available at http://cf.10xgenomics.com/supp/cell-exp/refdata-cellranger-mm10-3.0.0.tar.gz).

Samples were merged with the “cellranger aggregate” procedure without downsampling. Further analysis was performed starting from the UMI counts matrix by using the dropletUtils (version 1.5.4), scran (version 1.12), and scater (version 1.12)^38^ Bioconductor packages, following most steps illustrated in the simpleSingleCell Bioconductor workflow^39^.

Based on the clearly bimodal distributions observed across cells, cells were filtered out if they had log_10_ library sizes less than 3 (i.e., a minimum of 1,000 UMIs per library), log_10_ total number of features detected less than 2.5 (i.e. a minimum of 317 genes detected), or if they had 0%, or more than 7% of UMI counts attributed to the mitochondrial genes^40^. Low-abundance genes with average log_2_ counts per million reads lower than 0.002 were filtered out. The resulting filtered dataset included expression values for 12,182 genes for 943 cells, ranging from 189 to 334 cells per sample, for a total of 420 PBS cells, and 523 DEP cells. An average of 1,336 genes was detected per cell.

The expression values (log-transformed UMI counts) were normalized with library size factors estimated from pools of cells determined based on rank correlations cross expression profiles (scran function *quickCluster*)^39,41^. The technical noise was assumed to follow a Poisson distribution, and a mean-variance trend was fitted to the data (*makeTechTrend* function of the scran package with default parameters). This trend was subtracted to the variance of each gene to obtain the residual “biological” component of the variance. After performing a principal component analysis (PCA) on the top 500 most variable genes, the *denoisePCA* function of the scran package was used to choose the number of dimensions to retain in order to denoise the expression matrix.

Clustering cells into putative subpopulations was done on normalized and denoised log-counts values by using hierarchical clustering on the Euclidean distances between cells (with Ward’s criterion to minimize the total variance within each cluster; package cluster version 2.0.9). The cell clusters were identified by applying a dynamic tree cut (package dynamicTreeCut, version 1.63-1), which resulted in 4 clusters. The R package SingleR was used for reference-based annotation of the cell type of cells in our dataset^42^. We used the Immunological Genome Project (ImmGen) mouse database as reference^43^, and we filtered out the 31 cells not annotated as “Monocytes” or “Macrophages”, as these were likely contaminants.

Differential expression between DEP and PBS cells stratified by differentiation stage was performed by using a pseudo-bulk approach^44^. The UMI counts of cells from each sample and each cluster were aggregated. Cells from cluster 4 were ignored because they most likely represented damaged or low-quality cells (e.g. they displayed no expression of specific marker genes and had a higher fraction of reads coming from mitochondrial genes). Additionally, an “overall” analysis was performed where UMI counts of all cells from each sample were pooled. The resulting 16 pseudo-bulk samples were then treated as bulk RNA-seq samples for differential expression analysis. Genes were filtered to keep those with counts per million reads sequenced values higher than 1 in at least 2 samples, and detected in at least 5% of the cells of the cluster considered. The package edgeR (version 3.24.3)^45^ was used to perform TMM normalization^46^, and to test for differential expression with the Generalized Linear Model (GLM) framework. Genes with a false discovery rate (Benjamini-Hochberg method) lower than 5% were considered differentially expressed. Gene set enrichment analysis was performed with the function camera^47^ by using the default parameter value of 0.01 for the correlations of genes within gene sets, on gene sets from the Hallmark collection^48^ of the Molecular Signature Database (MSigDB, version 6.0)^49^. We considered only sets containing more than 5 genes, and gene sets with a false discovery rate lower than 20% were considered significant. Remaining statistical analysis on the expression dataset analysis and plotting were performed with the R software package (version 3.6.0).

### Data availability

The scRNA-seq dataset is available in the GEO database under accession GSE133406.

### Primary colon crypt cultures

Colon of untreated C57Bl/6N mice were isolated, washed, and cut into small pieces. The tissue was digested using 0.3 mg/mL collagenase XI (Sigma). To remove debris, the first two digestion steps (each 5 minutes) were discarded, followed by 3 x 10 minutes of digestion and cell collection. The collected cells were pooled, washed, and uniformly distributed on a 24 well plate previously coated with 0.1% gelatin (Sigma). The cells were allowed to settle overnight in an incubator at 37°C and 5% CO_2_. To prepare the cells for GLP-1 secretion, the cells were pre-incubated with Krebs Ringer solution (0.1 mM glucose) at 37°C for 15 minutes. To collect GLP-1 release, the Krebs Ringer solution was replaced with fresh Krebs Ringer solution (either 0.1 mM or 11.1 mM glucose) for 2 hours at 37°C with concomitant treatment with 125 μg/mL DEP or PBS as control. For normalization, the BCA of the cells was determined by using a Pierce BCA protein assay kit (Thermo Scientific).

### GSIS

Mouse islets were isolated by collagenase digestion (1.5mg/mL Colagenase IV) and subsequently purified by filtration and hand-picking. Islets were cultured free-floating in RPMI-1640 medium containing 11 mmol/l glucose and 10% FCS overnight, and were then washed and incubated in Krebs-Ringer buffer containing 2.8 or 20.0 mmol/l glucose and 0.5% BSA for 1 hour. Islet insulin was extracted with 0.18 mol/l HCl in 70% ethanol to determine insulin content. Secreted insulin and insulin content were assayed by a MSD insulin assay.

### Pharmacological depletion of macrophages

To pharmacologically deplete macrophages, mice were fed a diet containing the CSF1R-inhibitor PLX5622 (1200 ppm) or a control diet (Research Diets, New Brunswick, NJ, USA).

### IL-1β antibody treatment

Mice were exposed to DEP or PBS by oral gavage as described above. After 4 months of exposure, mice received a single injection of 10 mg/kg IL-1β antibody (01BSUR; Novartis) or isotype control (anti-cyclosporin A, Novartis). Treatment with the antibody was repeated after one week. From week 3 on, the mice received 5mg/kg once a week.

### Statistical analysis

Data are expressed as mean±SEM. The unpaired Mann-Whitney U test was used for statistical significance (GraphPad Prism). A p-value lower than 0.05 was considered statistically significant.

## Acknowledgments

We thank L. Rachid for technical assistance, M. Böni-Schnetzler/M. Donath for providing IL-1beta−/− mice, F. Cassee and L. Müller for their help with the particle procurement and dose estimation, M. Geiser for particle characterization and the members of the Donath, Hess, Recher, Berger and Mehling laboratories for advice and feedback. Single-cell RNA-sequencing was performed at the Ge-nomics Facility Basel of the ETH Zurich. Calculations were performed at sciCORE (http://scicore.unibas.ch/) scientific computing center at University of Basel. This study was support-ed by grants from the Velux Foundation, the Swiss Diabetes Foundation, the Wolfermann-Nägeli-Foundation, the Vontobel Foundation, and the Young Independent Investigator Research Grant from the Swiss Society for Endocrinology and Diabetology (all to CCW).

## Author contributions

Experimental design: AB, DM, CCW. Experimental execution: AB, TR, SAA, ZB, MS, TD, SM, JR, DM. Data analyses: AB, MS, JR, CCW. Figure preparation, manuscript writing: AB, CCW. Editing: AB, TR, SAA, ZB, JR, DM, CCW. CCW is the guarantor of this work.

## Author information

The authors declare no conflict of interest.

