## Extended data for "Air pollution mediates diabetes by disrupting gut innate mucosal immunity"

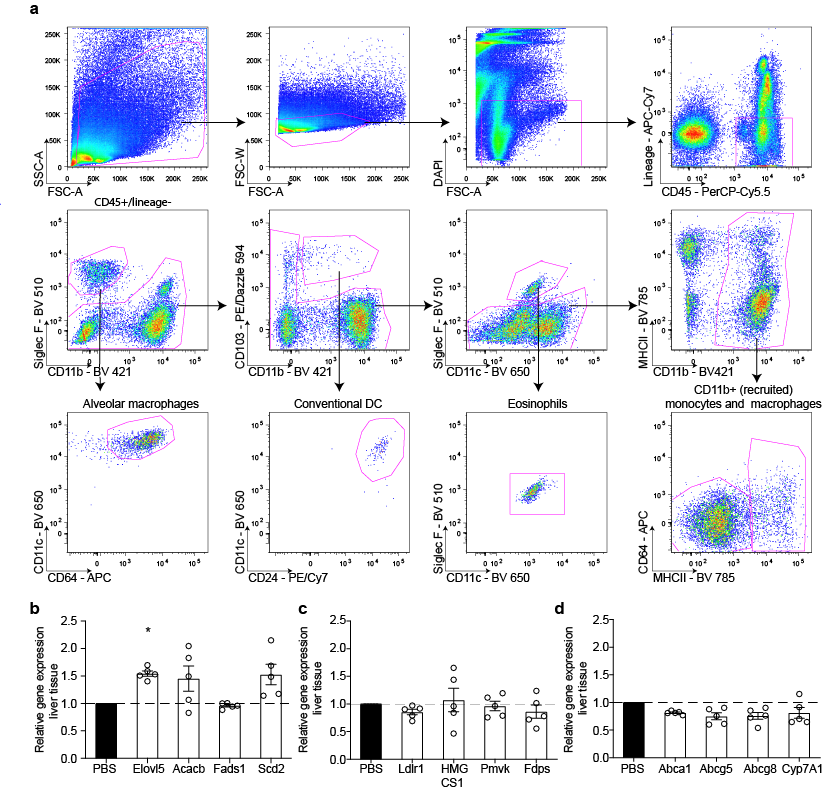

**Extended Data Fig. 1. Gating strategy of lung immune cells and gene expression of liver fatty acid synthesis, cholesterol synthesis, and lipid metabolism with exclusive lung exposure.** Gating strategy of lung immune cells (**a**). Gene expression of liver fatty acid synthesis (**b**), cholesterol synthesis (**c**), and lipid metabolism (**d**) in mice with exclusive lung exposure to DEP for 6 months compared to mice exposed to PBS. Data are presented as mean±SEM of 5 mice per group from one representative experiment compared by a two-tailed, unpaired Mann-Whitney U test (*p<0.05).

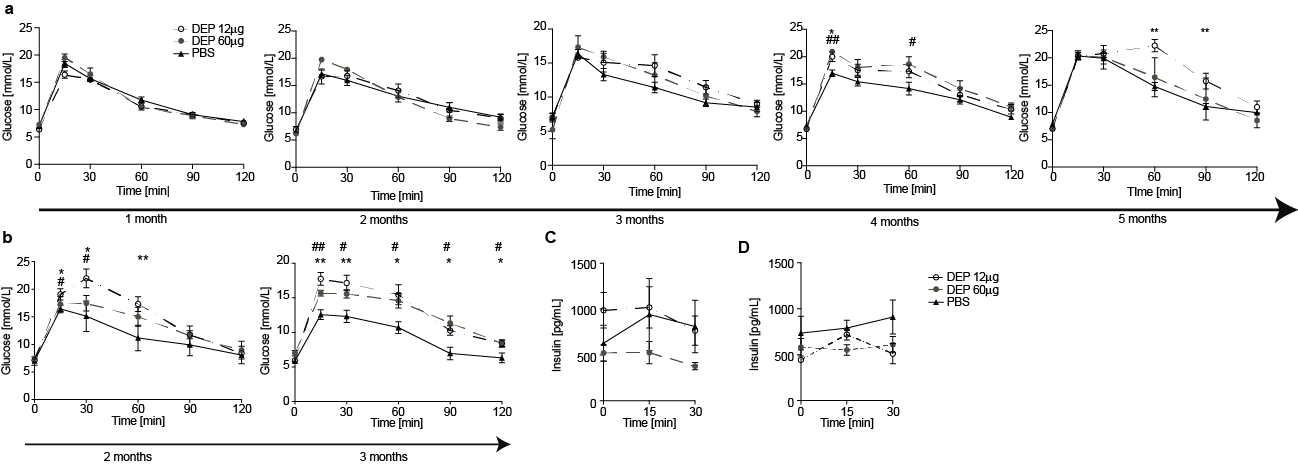

**Extended Data Fig. 2. Effect of oral air pollution exposure on glucose metabolism is independent of the dose.** Time course of glucose tolerance tests in mice exposed to either 12 or 60µg DEP or PBS via oral gavage during 5 months (**a**) and 3 months (**b**). Insulin after 5 months (**c**) and 3 months of exposure (**d**). Data are presented as mean±SEM of 3-5 mice per group from one representative experiment each compared by a two-tailed, unpaired Mann-Whitney U test (*p<0.05, **p<0.01). * indicates significances between 12µg DEP and PBS controls, # indicates significances between 60µg DEP and PBS controls.

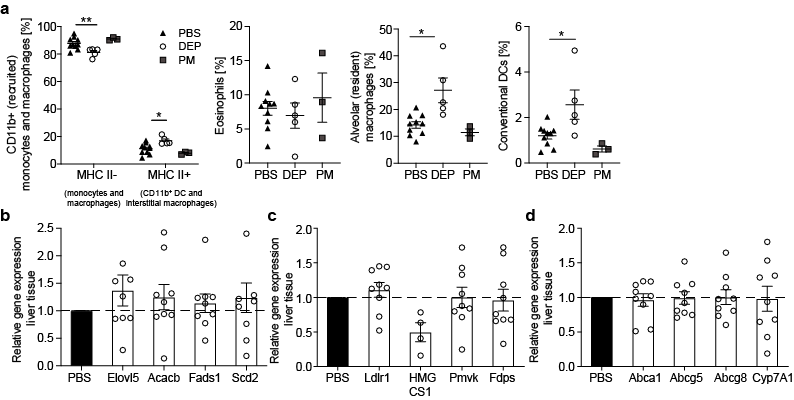

**Extended Data Fig. 3. Exclusive gut exposure to air pollutants does not induce lung inflammation or changes in liver fatty acid synthesis, cholesterol synthesis, and lipid metabolism.** Frequencies of recruited monocytes and macrophages among CD11b^+^ cells, eosinophils, alveolar macrophages, and conventional DCs gated on CD45^+^lineage^-^ cells in the lungs of mice orally exposed to DEP, PM, or PBS (**a**). Gene expression of liver fatty acid synthesis (**b**), cholesterol synthesis (**c**), and lipid metabolism (**d**) in mice with exclusive gut exposure to DEP for 6 months compared to mice exposed to PBS. Data are presented as mean±SEM of 3-5 mice per group from one representative experiment compared by a two-tailed, unpaired Mann-Whitney U test (*p<0.05, **p<0.01).

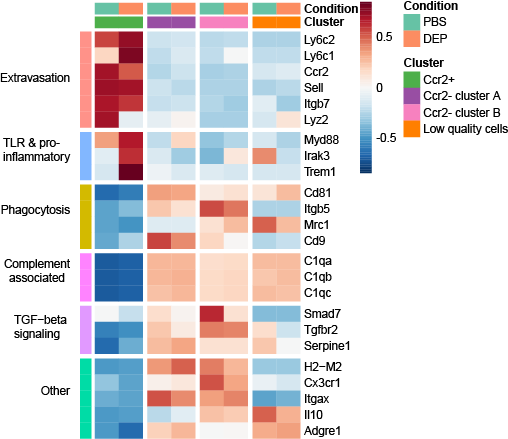

**Extended Data Fig. 4. Functionally defined genes in *Ccr2*^+^ and *Ccr2*^-^ intestinal macrophages.** Functionally defined subsets of mRNA transcripts that are up- or downregulated during differentiation from *Ccr2*^+^ to *Ccr2*^-^ intestinal macrophages of mice exposed to DEP or PBS. Data were obtained from one experiment with a group size of 2 animals per group.

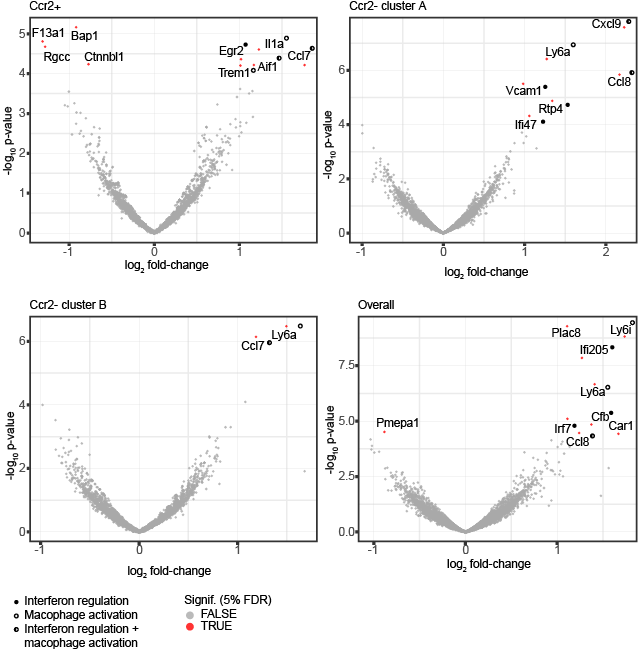

**Extended Data Fig. 5. Differences in genes expression in mice exposed to DEP or PBS.** Volcano plots comparing the expression of genes, which show a greater than two-fold change in expression of FACS sorted intestinal macrophages between mice exposed to DEP or PBS depicted by the different clusters (closed circles indicate genes related to interferon regulation, open circles genes related to macrophage activation, half-open circles to both). Data were obtained from one experiment with a group size of 2 animals per group.

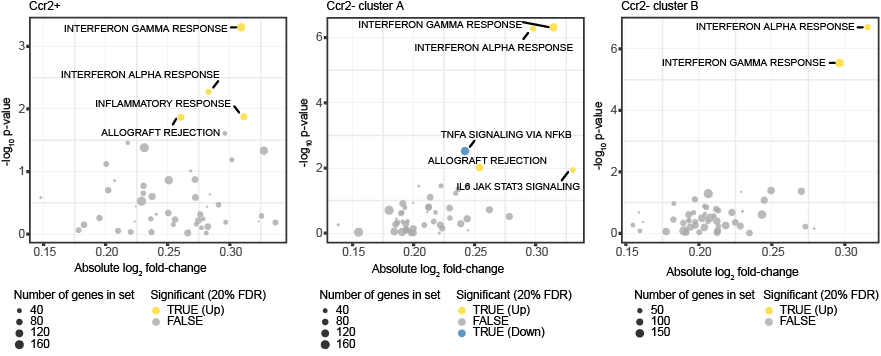

**Extended Data Fig. 6. Effects of DEP on MSigDB Hallmark pathways in *Ccr2*^+^ and *Ccr2*^-^ intestinal macrophages.** MSigDB Hallmark pathways that were activated (yellow) or repressed (blue) in intestinal macrophages upon exposure to DEP depicted for *Ccr2*^+^ and *Ccr2*^-^ clusters. Data were obtained from one experiment with a group size of 2 animals per group.

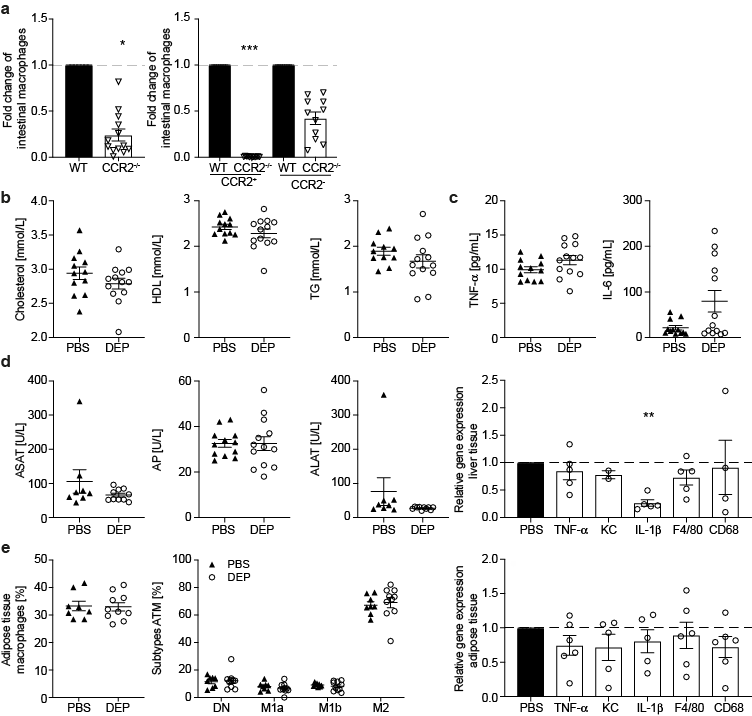

**Extended Data Fig. 7. CCR2-/- mice have low numbers of intestinal macrophages and no signs of dyslipidemia, systemic, adipose tissue, and liver inflammation upon oral exposure to DEP.** CCR2-/- mice were orally exposed to DEP or PBS for 6 months. Fold change of total, CCR2^+^ inflammatory and CCR2^-^ anti-inflammatory/resident macrophages in CCR2-/- mice compared to wild-type mice (**a**). Cholesterol, HDL, and TG (**b**), plasma TNF-α and IL-6 (**c**), liver lipids, liver enzymes and inflammatory gene expression in the liver (**d**). (**e**) Frequencies of macrophages in the adipose tissue among CD45^+^ cells, relative distribution of ATM subtypes, and inflammatory gene expression in adipose tissue of CCR2^-/-^ mice exposed to DEP compared to control mice exposed to PBS. Data are shown as mean±SEM of pooled data from 2-3 independent experiments compared by a two-tailed, unpaired Mann-Whitney U test (*p<0.05, **p<0.01, ***p<0.001).

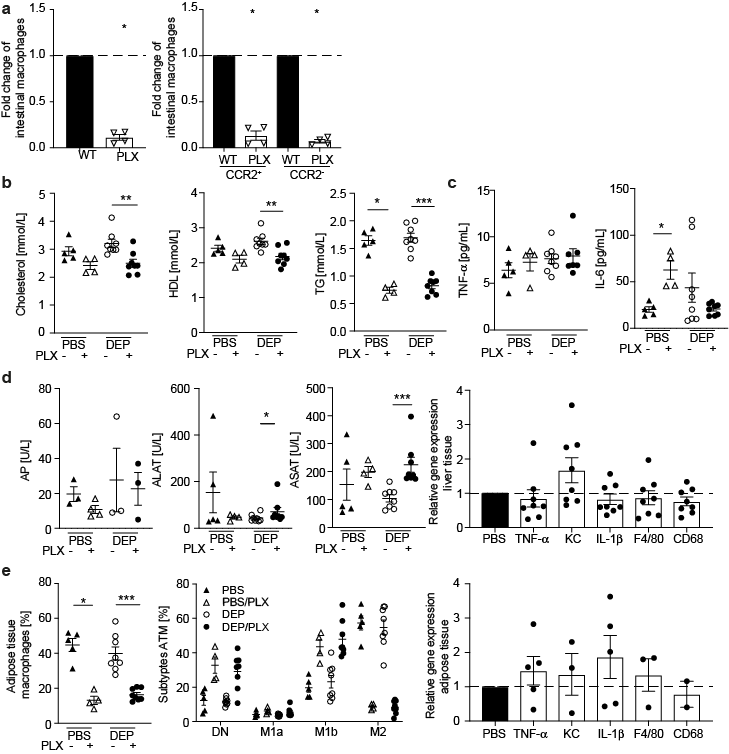

**Extended Data Fig. 8. Mice treated with CSF1R-inhibitor have low numbers of intestinal macrophages and no signs of dyslipidemia, systemic, adipose tissue, and liver inflammation upon oral exposure to DEP.** Mice were treated with CSF1R-inhibitor (PLX5622; PLX) to pharmacologically deplete macrophages and exposed to DEP or PBS for up to 10 months. Fold change of total, CCR2^+^ inflammatory and CCR2^-^ anti-inflammatory/resident macrophages in mice treated with PLX5622 compared to wild-type controls (**a**). Cholesterol, HDL and TG (**b**), plasma TNF-α and IL-6 (**c**), liver lipids, liver enzymes and inflammatory gene expression in the liver (**d**). (**e**) Frequencies of macrophages in adipose tissue among CD45^+^ cells, distribution of ATM subtypes and gene expression. Data are shown as mean±SEM of pooled data from 2 independent experiments compared by a two-tailed, unpaired Mann-Whitney U test (*p<0.05, **p<0.01, ***p<0.001).

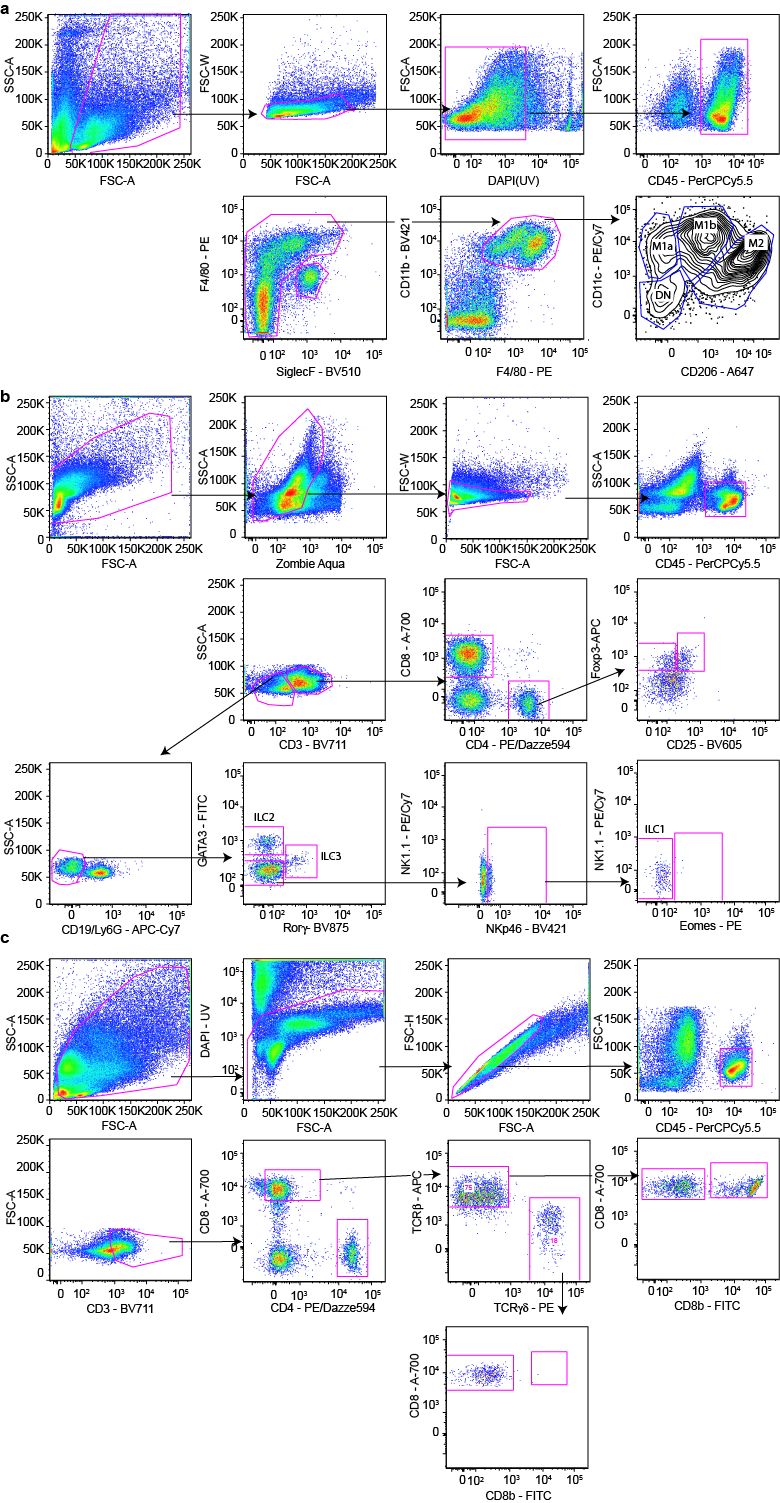

**Extended Data Fig. 9. Gating strategies.** (**a**) Adipose tissue cells were gated for live cells, followed by doublet exclusion with FSC W and A, and exclusion of dead cells by DAPI. CD45^+^ cells were then gated by expression of F4/80 and SiglecF. Eosinophils were defined as SiglecF^+^ F4/80^low^. CD11b and F4/80 double positive cells were further analyzed for the expression of CD206 and CD11b. (**b**) Colon lymphocytes were gated for live cells, and doublets excluded by FSC W and A. CD45^+^ cells were analyzed for T-cells and ILCs. T-cells were defined as CD3^+^ and either CD8^+^ and CD4^+^, the latter ones were further analyzed for Tregs defined as Foxp3^+^ and CD25^+^. ILCs were defied as CD3^-^/lineage^-^ cells, different subpopulations were further defined as Rorγ^+^ (ILC3), GATA3^+^ (ILC2) and Nkp46^+^ (ILC1). (**c**) IELs were gated for live cells, followed by doublet exclusion with FSC W and A, and exclusion of dead cells by DAPI. CD45^+^ cells were gated for CD3^+^ cells and further divided into CD4^+^ and CD8^+^ T-cells, CD8^+^ T-cells were further gated for expression of TCRαβ, TCRγδ, CD8αα, and CD8αβ.

Extended Data Table 1. Primers sequences used for quantitative real time-PCR

| **Gene** | **Forward Primer** | **Reverse Primer** |
| --- | --- | --- |
| **Housekeeping genes** | | |
| **B2m** | 5′ TTCTGGTGCTTGTCTCACTGA | 5′ CAGTATGTTCGGCTTCCCATTC |
| **Ppia** | 5′ GAGCTGTTTGCAGACAAAGTTC | 5′ CCCTGGCACATGAATCCTGG |
| **Inflammation markers** | | |
| **TNF-α** | 5′ ACTGAACTTCGGGGTGATCG | 5′ TGAGGGTCTGGGCCATAGAA |
| **IL-6** | 5′ GGATACCACTCCCAACAGACCT | 5′ GCCATTGCACAACTCTTTTCTC |
| **IL-1β** | 5′ GCAACTGTTCCTGAACTCAACT | 5′ ATCTTTTGGGGTCCGTCAACT |
| **KC** | 5′ CTGGGATTCACCTCAAGAACATC | 5′ CAGGGTCAAGGCAAGCCTC |
| **IL-10** | 5′ AGGCGCTGTCATCGATTTCTC | 5′ GCCTTGTAGACACCTTGGTCTT |
| **IL-18** | 5′TCTTGCGTCAACTTCAAGGA | 5′GTGAAGTCGGCCAAAGTTGT |
| **IL-22** | 5′TTG AGG TGT CCA ACT TCC AGC A | 5′AGC CGG ACG TCT GTG TTG TTA |
| **TGFβ_1_** | 5′CTCTCCACCTGCAAGACCAT | 5′CGAGCCTTAGTTTGGACAGG |
| **TGFβ_2_** | 5′GAAATACGCCCAAGATCGAA | 5′TGTCACCGTGATTTTCGTGT |
| **IFNγ** | 5′GTCTCTTCTTGGATATCTGGAGGAACT | 5′GTAGTAATCAGGTGTGATTCAATGACGC |
| **IL-17** | 5′ATC AGG ACG CGC AAA CAT GA | 5′TTG GAC ACG CTG AGC TTT GA |
| **Immune cells** | | |
| **CD68** | 5′ GCAGCACAGTGGACATTCAT | 5′ AGAGAAACATGGCCC GAAGT |
| **Adgre1 (Emr1)** | 5′ GCC CAG GAGTGGAATGTCAA | 5′ CAGACACTCATCAACATCTGCG |
| **Foxp3** | 5′ ACT CGC ATG TTC GCC TAC TTC AG | 5′ GGC GGA TGG CAT TCT TCC AGG T |
| **CD45** | 5′ATG GTC CTC TGA ATA AAG CCC A | 5′TCA GCA CTA TTG GTA GGC TCC |
| **CD3** | 5′ATG CGG TGG AAC ACT TTC TGG | 5′GCA CGT CAA CTC TAC ACT GGT |
| **CD20** | 5′ AACCTGCTCCAAAAGTGAACC | 5′ CCCAGGGTAATATGGAAGAGGC |
| **Regeneration markers islets** | | |
| **Ki67** | 5′ATC ATT GAC CGC TCC TTT AGG T | 5′GCT CGC CTT GAT GGT TCC T |
| **Ccna2a** | 5′ACA TTC ACA CGT ACC TTA GGG A | 5′CAT AGC AGC CGT GCC TAC A |
| **Cyclin D1** | 5′GCG TAC CCT GAC ACC AAT CTC | 5′CTC CTC TTC GCA CTT CTG CTC |
| **E2F1** | 5′TAG CCC TGG GAA GAC CTC AT | 5′CCC CAA AGT CAC AGT CAA AGA G |
| **Beta-cell identity** | | |
| **Pdx1** | 5′CCC CAG TTT ACA AGC TCG CT | 5′CTC GGT TCC ATT CGG GAA AGG |
| **Foxo1** | 5′GTA CGC CGA CCT CAT CAC CA | 5′TGC TGT CGC CCT TAT CCT TG |
| **Ins2** | 5′CCC TGC TGG CCC TGC TCT T | 5′AGG TCT GAA GGT CAC CTG CT |
| **LXR** | 5′TCT GGA GAC GTC ACG GAG GTA | 5′CCC GGT TGT AAC TGA AGT CCT T |
| **Abca1** | 5′GGA CAT GCA CAA GGT CCT GA | 5′CAG AAA ATC CTG GAG CTT CAA A |
| **Abcg5** | 5′TGG ATC CAA CAC CTC TAT GCT AAA | 5′GGC AGG TTT TCT CGA TGA ACT G |
| **Abcg8** | 5′TGC CCA CCT TCC ACA TGT C | 5′ATG AAG CCG GCA GTA AGG TAG A |
| **Cyp7A1** | 5′AGC AAC TAA ACA ACC TGC CAG TAC TA | 5′GTC CGG ATA TTC AAG GAT GCA |
| **Fatty acid synthesis** | | |
| **Elovl5** | 5′ CTGAGTGACGCATCGAAATG | 5′ CTTGCACATCCTCCTGCTC |
| **Acacb** | 5′ CCCAGGAGGCTGCATTGA | 5′ AGACATGCTGGGCCTCATAGTA |
| **Fads1** | 5′ TGGTGCCCTTCATCCTCTGT | 5′ GGTGCCCAAAGTCATGCTGTA |
| **Scd2** | 5′ TGCCTTGTATGTTCTGTGGC | 5′ TCCTGCAAGCTCTACACCTG |
| **Cholesterol pathway** | | |
| **Ldlr1** | 5′ ACCTGCCGACCTGATGAATTC | 5′ GCAGTCATGTTCACGGTCACA |
| **HmgCS1** | 5′ TTTGATGCAGCTGTTTGAGG | 5′ CCACCTGTAGGTCTGGCATT |
| **Pmvk** | 5′ GCTCGCATCCAGAAGTCTCT | 5′ GCTCTCTGGTCCACTCAAGG |
| **Fdps** | 5′ GAGTCTGCCCGATCTCTGTC | 5′ TGAACCTGCTGGAGCTCTTT |
| **LXR-target genes** | | |
| **Abca1** | 5′ GGACATGCACAAGGTCCTGA | 5′ CAGAAAATCCTGGAGCTTCAAA |
| **Abcg5** | 5′ TGGATCCAACACCTCTATGCTAAA | 5′ GGCAGGTTTTCTCGATGAACTG |
| **Abcg8** | 5′ TGCCCACCTTCCACATGTC | 5′ ATGAAGCCGGCAGTAAGGTAGA |
| **Cyp7A1** | 5′ AGCAACTAAACAACCTGCCAGTACTA | 5′ GTCCGGATATTCAAGGATGCA |
